## Supplementary Methods, Tables and Figures for "Genome Methylation Predicts Age and Longevity of Bats"

This supplement contains the following text, tables and figures.

**Supplementary Methods**

**Supplementary References**

**Supplementary Table 1. Bat genome assemblies used for identifying CpG sites.**

**Supplementary Table 2. PGLS model coefficients for predicting log(maximum lifespan) by log mass and mean rate of methylation.**

**Supplementary Fig. 1. Epigenetic age predictions for 12 species of bats.**

**Supplementary Fig. 2. Combined epigenetic age predictions for all species and for three genera of bats.**

**Supplementary Fig. 3. Longevity differentially methylated positions (DMPs) exhibit more rapid change in DNAm with age for short-lived than long-lived bat species.**

**Supplementary Fig. 4. Genome positions of probes on the mammalian methylation array are conserved in bats.**

**Supplementary Fig. 5. Differentially methylated positions (DMPs) in three longlived bats exhibit very similar overlap, genome location, and gene associations.**

**Supplementary Fig. 6. Age and longevity DMPs are near genes associated with immunity and cancer in long-lived bats.**

**Supplementary Methods**

The number of tissue samples per species and per sex, as well as the range of ages of individuals of each bat species, are provided in Table 1 of the Methods. Here, we provide additional information on when and where samples were taken from either captive or free-ranging animals for each of the 26 species of bats used in this study.

Pallid bats, *Antrozous pallidus,* were captured between 2005 and 2008 at six sites in central Oregon (44.94° N, 120.38° W) using mist nets over a water source or outside a night roost or with a handnet on an extension pole outside a day-roosting crevice. Each bat was weighed, measured and marked with a numbered band. Adults were distinguished from juveniles by closed epiphyseal gaps[1]. Tissue samples were obtained from wing membranes using 3 mm biopsy punches and stored in 95% ethanol until DNA was extracted using a Qiagen DNeasy Tissue Kit. DNA extracts were stored frozen at -80°C. Live animal procedures conformed to the American Society of Mammalogists guidelines[2] and were approved by the University of Maryland Institutional Animal Care and Use Committee (protocol R-08-39).

Wing tissue from *Artibeus jamaicensis, Cynopterus brachyotis, Eidolon helvum, Pteropus giganteus, P. hypomelanus, P. poliocephalus, P. pumilus, P. rodricensis, P. vampyrus, and Rousettus aegyptiacus* was taken from bats kept at the Lubee Bat Conservancy, a USDA registered facility, in Gainesville, Florida. The bats are group-housed in twelve 1068 sq. ft. enclosures with indoor temperature-controlled roosting areas and outdoor flight rooms and are fed a diet of fruit, vegetables and nutritional supplements. Wing tissue biopsies are periodically taken from individually marked animals and kept at -20°C in 95% ethanol. The majority of 243 samples from these species were taken from animals that were born in captivity. DNA was extracted with a Zymo miniprep plus kit.

Wing tissue samples were taken from captive *Carollia perspicillata* housed in a tropical zoo (Papiliorama, Kerzers FR, Switzerland). Approximately 400 bats roost in an artificial cave kept on a reversed light cycle and are fed twice a night with a fruit-based diet. Since 2011, the population has been monitored by capturing individuals using a harp-trap placed at the entrance to the cave. Forearm length, body weight, reproductive status and tooth-wear are recorded from every captured individual. At first capture, individuals are marked on the forearms with a unique combination of three colored plastic rings (A.C. Hughes, UK, size XB). Between July and November 2018, 3mm biopsies were punched on the patagium and hermetically stored in silica gel. Based on the date of first capture and tooth-wear score, the age of each individual sampled was estimated[3]. DNA was extracted with a Zymo miniprep plus kit.

Biopsy punches (2 or 3 mm) were taken from the wing of captive common vampire bats, *Desmodus rotundus,* between 2010 and 2014for husbandry details see [4]. Bats were housed in flight cages (3 x 2 x 1.5 m) as a captive group at the Cranbrook Institute of Science (24-39 bats, Bloomfield Hills, MI, USA) or at the University of Maryland (7 bats, University of Maryland Institutional Animal Care and Use Committee protocol R-10–63). Age was determined based on zoo birth records. Individuals were born at the Houston Zoo, Cincinnati Zoo, Chicago Brookfield Zoo, or the Cranbrook Institute of Science. Tissue samples were stored in 95% ethanol prior to DNA extraction using a Qiagen DNeasy kit. DNA extracts were frozen for long-term storage at -80°C. Live animal procedures conformed to the American Society of Mammalogists guidelines and was approved by the University of Maryland Institutional Animal Care and Use Committee (protocol R-10-63).

Wing tissues were sampled in August 2018 and 2019 with a 3 mm biopsy punch from the wing of captive big brown bats (*Eptesicus fuscus*) of known age[5,6]. The Cooper Laboratory at

Northeast Ohio Medical University (NEOMED; Rootstown, Ohio) maintains[7] a colony of known-aged big brown bats transferred from a colony previously maintained by Dr. Ellen Covey at the University of Washington. This colony was started in 2005 with bats caught in North Carolina and were banded according to year of capture or birth. These bats underwent natural hibernation and were exclusively fed an *ab libitum* diet of fresh water and mealworms (*Tenebrio molitor*) of known nutrient content. In 2014 bats were transported to NEOMED and are now housed indoors on a 12 h light/dark cycle and fed the same *ab libitum* fresh water and mealworm diet. Tissue samples were stored frozen at -80°C in DNA Shield prior to DNA extraction with a Zymo miniprep plus kit.

During July 2019, wing membrane samples were obtained from subadult or adult female and subadult male lesser long-nosed bats, *Leptonycteris yerbabuenae,* with a 4 mm biopsy punch at the entrance of the Pinacate Cave in the Reserva de la Biosfera el Pinacate y Gran Desierto de Altar (31°38'51.6" N, 113°28'53.5" W), Sonora, Mexico. Bats were captured using mist nets (Avinet models: TB02, TB06, TB012; Portland, Maine, USA) set outside caves just prior to when bats emerged to forage. Individuals were sexed, weighed and the forearm measured. To discriminate subadults from adults, age was determined by the degree of fusion of the epiphyses at the metacarpal–phalangeal joint. Tissue samples were stored in DNA/RNA Shield buffer (Zymo Scientific, Irvine, CA 92614, U.S.A.). DNA was extracted with a Zymo miniprep plus kit. Bat tissue samples were collected under permit SGPA/DGVS/06361/17 issued to R. A. Medellín by The Ministry of Environment and Natural Resources.

Samples of velvet free-tailed bats, *Molossus molossus*, come from a long-term study[8,9,10] in Gamboa, Panama (09°07’ N 79°41’ W), where the bats roost in crevices in houses. We captured social groups with mist nets (Ecotone, Gydnia, Poland) at the entrance of roosts during evening emergence and individually marked all bats with a subcutaneous passive integrated transponder (Trovan ID-100, Euro ID, Weilerswist, Germany) at first capture. Wing tissue samples were taken with a 3 mm biopsy punch and stored in 96% ethanol until DNA extraction using a Zymo miniprep kit. Capture and handling of animals were carried out under permits from the Autoridad Nacional del Ambiente in Panama with approval from the Institutional Animal Care and Use Committee of the Smithsonian Tropical Research Institute (2012-0505-2015).

Little brown bats, *Myotis lucifugus*, were captured as they departed from an attic maternity colony in Chestertown, Maryland, in September 1996. Captured bats were weighed, measured and banded with individually marked bands. Young of the year were identified by their weight and absence of tooth wear. Wing membrane biopsies were taken and stored in a 5M NaCl with 20% dimethyl sulfoxide solution and kept frozen at -80°C. DNA was extracted with a Zymo miniprep plus kit. Bat capture and handling was approved by the Maryland Department of Natural Resources.

Wing tissue samples were taken from greater mouse-eared bats, *Myotis myotis*, between 2013 and 2018 as part of a long-term mark-recapture study conducted by Bretagne Vivante in Brittany, France[11,12]. Bats were caught using modified harp traps as they left the roost. Individuals at first capture are fitted with PIT tags to facilitate identification on subsequent recaptures. Measurements taken from each individual include sex, forearm length, weight and transponder number. Age class (juvenile or adult) is determined by examining the degree of the epiphyseal closure of the metacarpal-phalangeal joints[13]. Wing biopsies were taken with a 3 mm biopsy punch, flash frozen and stored in liquid nitrogen prior to extraction. All procedures were conducted with full ethical approval and permission (AREC-13-38-Teeling) awarded by the University College Dublin ethics committee. DNA was extracted from wing biopsies using a

Promega Wizard SV DNA extraction kit (catalog no. A2371) or the Qiagen DNeasy Blood and Tissue kit (Qiagen). Extractions carried out with the Promega kit were partially automated using a Hamilton STAR Deck liquid handling robot.

Adult and juvenile Mexican fishing bats, *Myotis vivesi*, were captured by gloved hand from roosts in talus slopes on Isla Partida Norte in the Gulf of California, Mexico (29°03'N, 113°00'W) during the day between 2015 and 2018[14,15]. Individuals were measured and banded with numbered metal bands on their forearms for identification upon recapture. In 2018 wing tissue was taken with a 3 mm biopsy punch and preserved in Zymo DNA shield. Bat capture and handling were conducted under permits #7668–15 and 2492–17 from Dirección General de Vida Silvestre, and permits #17–16 and 21–17 from Secretaría de Gobernación, and the University of Maryland Institutional Animal Care and Use Committee protocols FR-15-10 and FR-18-20.

Common noctules, *Nyctalus noctula*, were captured as part of a long-term study[16,17,18] at the Seeburgpark in Kreuzlingen, Switzerland (47.649928° N, 9.186123° E) where bats regularly roost in boxes. Each bat was marked with a subcutaneous pit-tag (ID100; Euro ID, Weilerswist, Germany) injected under the dorsal skin. Wing tissue samples were taken with a 3 mm biopsy punch and stored in 96% ethanol until DNA extraction using a Zymo miniprep kit. All handling and sampling of the bats in Switzerland was approved by the Veterinäramt Thurgau (permit FIBL1/12).

Wing tissue samples were taken in September 2018 from lesser spear-nosed bats, *Phyllostomus discolor*, kept in a breeding colony in the Department Biology II of the Ludwig-Maximilians-University in Munich. In this colony animals were kept under semi-natural conditions (12 h day/night cycle, 65 to 70 % relative humidity, 28°C) with free access to food and water. The license to keep and breed *P. discolor* was issued by the German Regierung von

Oberbayern. Under German Law on Animal Protection a special ethical approval is not needed for wing tissue collection. Wing tissue was stored in RNAlater until DNA was extracted using QIAamp® MinElute columns following the manufacturer's instructions. The samples were eluted in 50µl of molecular grade water and concentrated to reduce their volume by approximately 50% using a Speedvac, with the following settings: duration 20 minutes, temperature in the chamber 30°C, $H_2O$ (water) mode.

Greater spear-nosed bats, *Phyllostomus hastatus,* were captured and sampled between 1990 and 2018 in Trinidad, Lesser Antilles[19,20,21,22]. Most often, harem groups, which include one adult male plus 15-20 lactating females with pups, were captured during the day from within a solution depression in the ceiling of either Tamana (10.4711°N, 61.1958°W), Caura (10.7019°N, 61.3614°W), or Guanapo cave (10.6942°N, 61.2654°W) using a bucket trap. Captured bats were sexed, measured for size, weight, and tooth wear, and individually marked with stainless steel numbered bands. Age was determined exactly for adults that were recaptured after being banded as pups. Wing biopsy punches (4 mm) were stored frozen at -80°C in either a 5M NaCl with 20% dimethyl sulfoxide solution or Zymo DNA Shield prior to DNA extraction using a Qiagen Puregene or Zymo miniprep plus kit. Frozen samples were selected to maximize the number of known-age individuals with approximately equal numbers at all ages. Animal handling methods follow guidelines by the American Society of Mammalogists and were approved by the University of Maryland Institutional Animal Care and Use Committee (protocols R-91-33, R-94-25, R-01-07, R-11-21, R-13-77) under licenses from the Forestry Division of the Ministry of Agriculture, Land and Fisheries, Trinidad and Tobago.

Wing tissue samples were taken from greater horseshoe bats, *Rhinolophus ferrumequinum*, by using 3 mm biopsy punches between 2016 and 2018 from wild female bats as part of a long-term study at a maternity colony in Gloucestershire, UK[23,24,25]. Bats were captured at the roost with hand nets, and all individuals were weighed, ringed with aluminum alloy rings and morphometric data such as forearm length recorded under licenses (Natural England Project Licences 2015-9918-SCI-SCI; 2016-23583-SCI-SCI; 2017-30137-SCI-SCI) issued to Roger Ransome. All bats studied were first marked as infants, so we could be certain of their age. Bats were aged between 1-21 years, with the 40 individuals selected in a fairly even manner across this age span. Sampling procedures were conducted under licenses (Natural England 2015-11974-SCI-SCI; 2016-25216-SCI-SCI; 2017-31148-SCI-SCI) issued to Gareth Jones, with tissue biopsy additionally licensed under Home Office Project Licenses (PPL 30/3025 prior to 2018; P307F1428 from 2018 onwards) and Home Office personal licences. Tissue samples were stored in silica gel beads and then transferred to a -20C freezer for long-term storage. DNA was extracted from wing biopsies using a Promega Wizard SV DNA extraction kit or a Qiagen DNeasy Blood and Tissue kit. Extractions carried out with the Promega kit were partially automated using a Hamilton STAR Deck liquid handling robot but otherwise followed manufacturer's instructions.

Samples of proboscis bats, *Rhynchonycteris naso,* came from a long-term study between 2005 and 2016 at La Selva Biological Station in Costa Rica (10° 25′ N, 84° 00′W) [86-88]. Bats were mist-netted in the vicinity of their roosts. Wing tissue was sampled with a 4 mm biopsy punch, individuals were marked with colored plastic bands, sexed, measured and age class determined (juvenile: 0-4 months, subadult: 5-10 months, or adult>10 months)[26]. Age was determined exactly for individuals that were banded as pups and recaptured as adults. Ethanol (80%) was used to preserve tissue samples, and a salt–chloroform procedure or Qiagen BioSprint 96 DNA Blood Kit was used for DNA isolation[26,27].

Samples of greater sac-winged bats, *Saccopteryx bilineata* came from long-term studies in Costa Rica (n = 21 from La Selva Biological Station, 10° 25′ N, 84° 00′W and n = 6 from Santa Rosa National Park, 10° 53′ N, 85° 46′ W) and Panama (n = 4 from the Biological Station Barro Colorado Island (BCI) of the Smithsonian Tropical Research Institute, 9° 9′ N/79° 51′ W) between 1994 and 2016[28,29]. Bats were captured with mist nets when entering or leaving their day roosts, individually banded with two coloured plastic bands on their forearms, and a wing tissue biopsy sample (4mm) preserved in 80% ethanol was taken. Age was determined exactly for individuals that were banded as pups and recaptured as adults. DNA was extracted with a salt-chloroform procedure or with the Qiagen BioSprint 96 DNA Blood Kit[28,30].

Mexican free-tailed bats, *Tadarida brasiliensis*, are housed at Bat World Sanctuary, a licensed non-profit bat rehabilitation facility and accredited sanctuary in Weatherford, Texas. Most individuals sampled were rescued as pups, although some were rescued as adults, making their exact age unknown. Individuals are group housed in large indoor enclosures. Wing membrane biopsies (4 mm) were collected in August 2019 and stored in Zymo DNA Shield until DNA was extracted using a Zymo miniprep plus kit.

**Supplementary Table 1.** Number of CpG sites mapped per bat genome assembly.

| Species | Assembly and annotation | Source | CpGs |
|---|---|---|---|
| *Molossus molossus* | HLmolMol2 | MPI* | 29596 |
| *Myotis myotis* | HLmyoMyo6 | MPI* | 28945 |
| *Phyllostomus discolor* | HLphyDis3 | MPI* | 29294 |
| *Rhinolophus ferrumequinum* | HLrhiFer5 | MPI* | 30724 |
| *Pipistrellus kuhlii* | HLpipKuh2 | MPI* | 27164 |
| *Rousettus aegyptiacus* | HLrouAeg4 | MPI* | 30576 |
| *Desmodus rotundus* | GCF 002940915.1, ASM294091v2 | NCBI | 28552 |
| *Eptesicus fuscus* | GCF 000308155.1, EptFus1.0 | NCBI | 28606 |
| *Myotis lucifugus* | GCF 000147115.1, Myoluc2.0 | NCBI | 26474 |
| *Pteropus vampyrus* | pteVam1.100 | ENSEMBL | 22216 |

MPI* (downloaded from https://bds.mpi-cbg.de/hillerlab/Bat1KPilotProject/)

**Supplementary Table 2.** PGLS model coefficients for predicting log(maximum lifespan).

| Methylation[1] | Variable | Coefficient | SE | t | P |
|---|---|---|---|---|---|
| Hyper | Intercept | 1.366 | 0.125 | 10.90 | 7.31E-10 |
| | Mean rate | -11.414 | 1.965 | -5.81 | 1.11E-05 |
| | Log mass | 0.134 | 0.060 | 2.23 | 3.75E-02 |
| Hypo | Intercept | 1.485 | 0.136 | 10.96 | 6.67E-10 |
| | Mean rate | -6.710 | 1.136 | -5.91 | 8.87E-06 |
| | Log mass | 0.047 | 0.062 | 0.76 | 4.58E-01 |

[1]Mean methylation rate is based on 1175 hypermethylating sites and 825 hypomethylating sites for 23 species, each with 10 or more known-aged individuals

**Supplementary Figure Legends**

**Supplementary Fig. 1. Epigenetic age predictions for 12 species of bats.** Each panel displays the results of separate leave-one-out (LOO) cross-validation estimates of age based on penalized regression of the DNAm values for each the 12 bat species with the largest sample sizes (N) not shown in the main text. Correlation between observed chronological age and predicted age and median absolute error (MAE) are indicated at the top of each panel.

**Supplementary Fig. 2. Combined epigenetic age predictions for all species and for three genera of bats.** a) Results of separate leave-one-out (LOO) cross-validation estimates of age based on penalized regression of the DNAm values for each of 23 colorcoded bat species. Species with fewer than 10 samples are omitted. b) LOO cross-validation estimate of age based on penalized regression of the DNAm values for six species of *Pteropus*. c) LOO cross-validation estimate of age based on penalized regression of the DNAm values for two species of *Phyllostomus*. d) LOO cross-validation estimate of age based on penalized regression of the DNAm values for three species of *Myotis*. Sample size (N), correlation between observed chronological age and predicted age, significance, and median absolute error (MAE) are indicated at the top of each panel.

**Supplementary Fig. 3. Longevity differentially methylated positions (DMPs) exhibit greater age-related change in DNAm for short-lived than long-lived bats.** DNAm plotted against squareroot (age+1) for three long-lived species (DERO = *Desmodus rotundus*, MYMY = *Myotis myotis,* RHFE = *Rhinolophus ferrumequinum*) and two short-lived species (LEYE =

*Leptonycteris yerbabuenae,* MOMO = *Molossus molossus*) for four probes where the longevity x age interaction was highly significant (P < 10e-5). Least squares regression lines displayed with shaded regions indicating confidence intervals for a) cg21759108 which is in an exon of CTNNA3 (catenin alpha 3), b) cg24840164 which is in an exon of SATB2 (SATB homeobox 2), c) cg09996908 which is in an intron of IGDCC3 (immunoglobulin superfamily DCC subclass member 3) and d) cg07279255 which is in the promoter region of EN1 (engrailed homeobox 1).

**Supplementary Fig. 4. Genome positions of probes on the mammalian methylation array are conserved in bats.** a) Probes that map to a bat genome are in similar locations relative to the transcription start site (TSS) for the species in Supplementary Table 1. Probes map to distal intergenic regions more often in bats than in humans. b) Most probes (~80%) map downstream of the TSS in all bats. Overrepresented regions include 10-100 kb downstream of TSS (40%), distal (>100kb downstream of TSS) intergenic region (20%), and promoter regions within 1kb of TSS (~15%). c) Probes were more likely to be near the same gene among bats (median = 93%) than in bat vs human comparisons (median = 86%). The proportion of probes nearest the annotated gene was calculated for each pair of bat species and human, with each comparison plotted as a point overlaying the boxplot. Probes not mapped in either species were not counted. d) Conservation of probe location is high between bats used for identifying longevity DMPs. Long-lived vs. long-lived refers to comparisons between any two of *R. ferrumequinum, M. myotis,* and *D. rotundu*s, long-lived vs short-lived refers to comparisons between *M. molossus* and any long-lived species, and “NA” or “Neither” refers to any other combination. Probes associated with Longevity and Age DMPs are most conserved.

**Supplementary Fig. 5. Differentially methylated positions (DMPs) in three longlived bats exhibit very similar overlap, gene associations, and genome location**. a) Age DMPs overlap 20% with hypermethylating (+) and hypomethylating (-) longevity DMPs in the three long-lived bat species used to identify longevity DMPs. b) Number of unique orthologous to human genes nearest age and longevity DMPs for the three long-lived bat species. Sign on numbers in overlap region indicate methylation direction for age then longevity. c) Age DMPs are highly enriched near promoter regions, i.e. 100 to -10,000 bp from the transcription start site, and over 95% exhibit hypermethylation in the three long-lived bat species. d) Longevity DMPs are also enriched in promoter regions with more than 80% exhibiting hypermethylation in the three long-lived bat species.

**Supplementary Fig. 6. Age and longevity DMPs are near genes associated with immunity and cancer in long-lived bats.** a) Gene ontology enrichment analysis of biological process for unique genes from promoter regions in the long-lived bat, *Rhinolophus ferrumequinum,* reveals significant enrichment only for hypermethylating age and longevity genes. Longevity genes are enriched for a subset of age-associated processes. The other two long-lived bat species show similar patterns. Only three significant GO terms from each parent-child group are shown to minimize redundancy. b) Enrichment analysis of protein class for unique *R. ferrumequinum* genes from promoter regions reveals significant enrichment of transcription factors (TF) in both age and longevity genes. The other two long-lived bat species show similar patterns. Cell color indicates significance (negative log P for GO terms with adjP < 10e-4) of enrichment in a) and b). c) Top nine transcription factors associated with age or longevity gene promoters in *R. ferrumequinum,* with integrative rank significance (see Methods) indicated as negative log P.

Only genes with hypermethylated sites in promoter regions were significantly enriched. d) Genes near DMPs associated with longevity overlap genes involved in innate immunity or cancer for each of the three long-lived bat species. For *R. ferrumequinum,* longevity genes are associated with cancer genes ($P = 0.001$, FET) and with immunity genes ($P < 0.001$, FET). In *D. rotundus,* longevity genes are marginally associated with cancer genes ($P = 0.063$, FET) but more strongly with immunity genes ($P = 0.017$, FET). For *M. myotis*, longevity genes are associated with cancer genes ($P = 0.030$, FET) and with immunity genes ($P = 0.025$, FET).

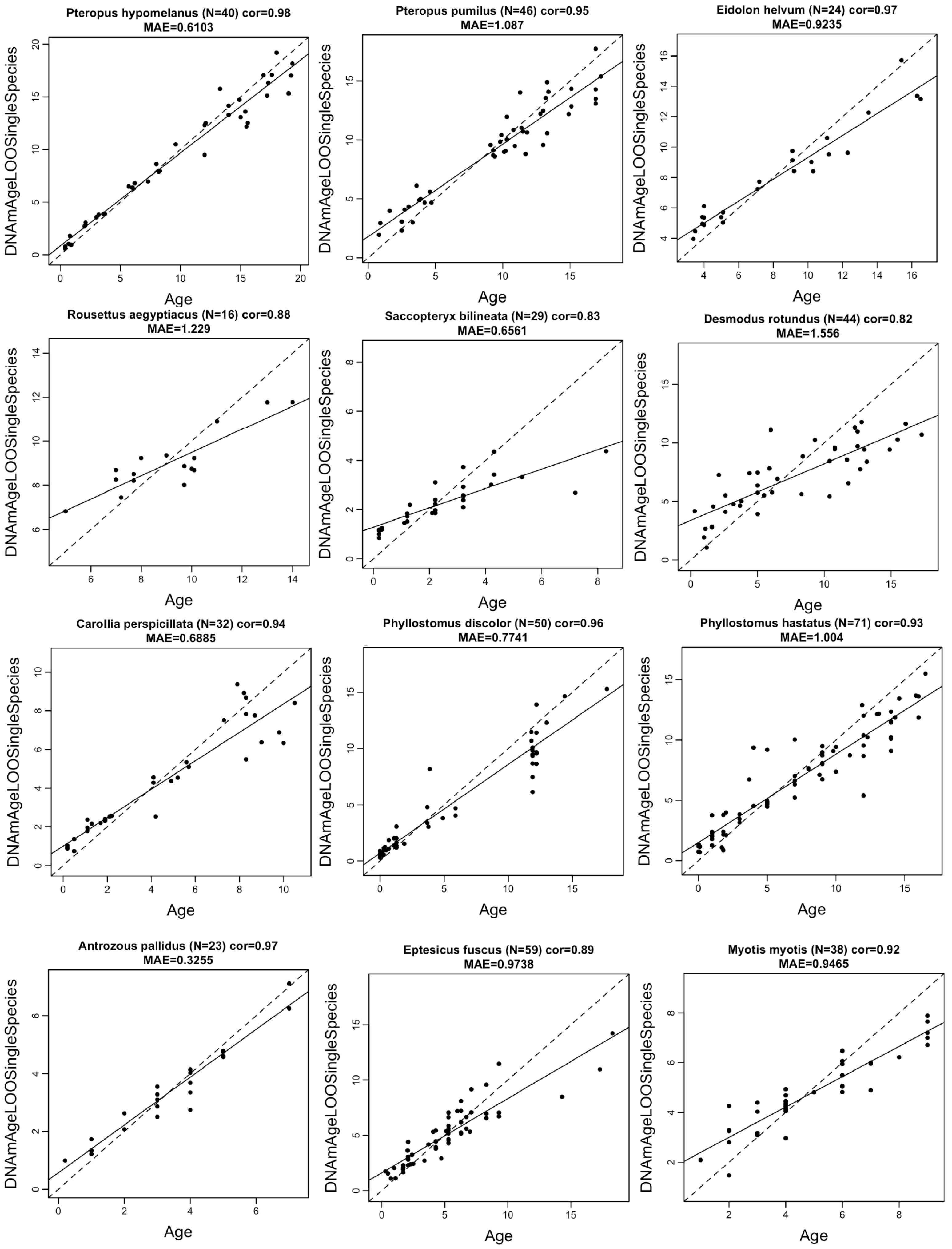
Pteropus hypomelanus (N=40) cor=0.98
MAE=0.6103
Pteropus pumilus (N=46) cor=0.95
MAE=1.087
Eidolon helvum (N=24) cor=0.97
MAE=0.9235
Rousettus aegyptiacus (N=16) cor=0.88
MAE=1.229
Saccopteryx bilineata (N=29) cor=0.83
MAE=0.6561
Desmodus rotundus (N=44) cor=0.82
MAE=1.556
Carollia perspicillata (N=32) cor=0.94
MAE=0.6885
Phyllostomus discolor (N=50) cor=0.96
MAE=0.7741
Phyllostomus hastatus (N=71) cor=0.93
MAE=1.004
Antrozous pallidus (N=23) cor=0.97
MAE=0.3255
Eptesicus fuscus (N=59) cor=0.89
MAE=0.9738
Myotis myotis (N=38) cor=0.92
MAE=0.9465
DNAmAgeLOOSingleSpecies
Age

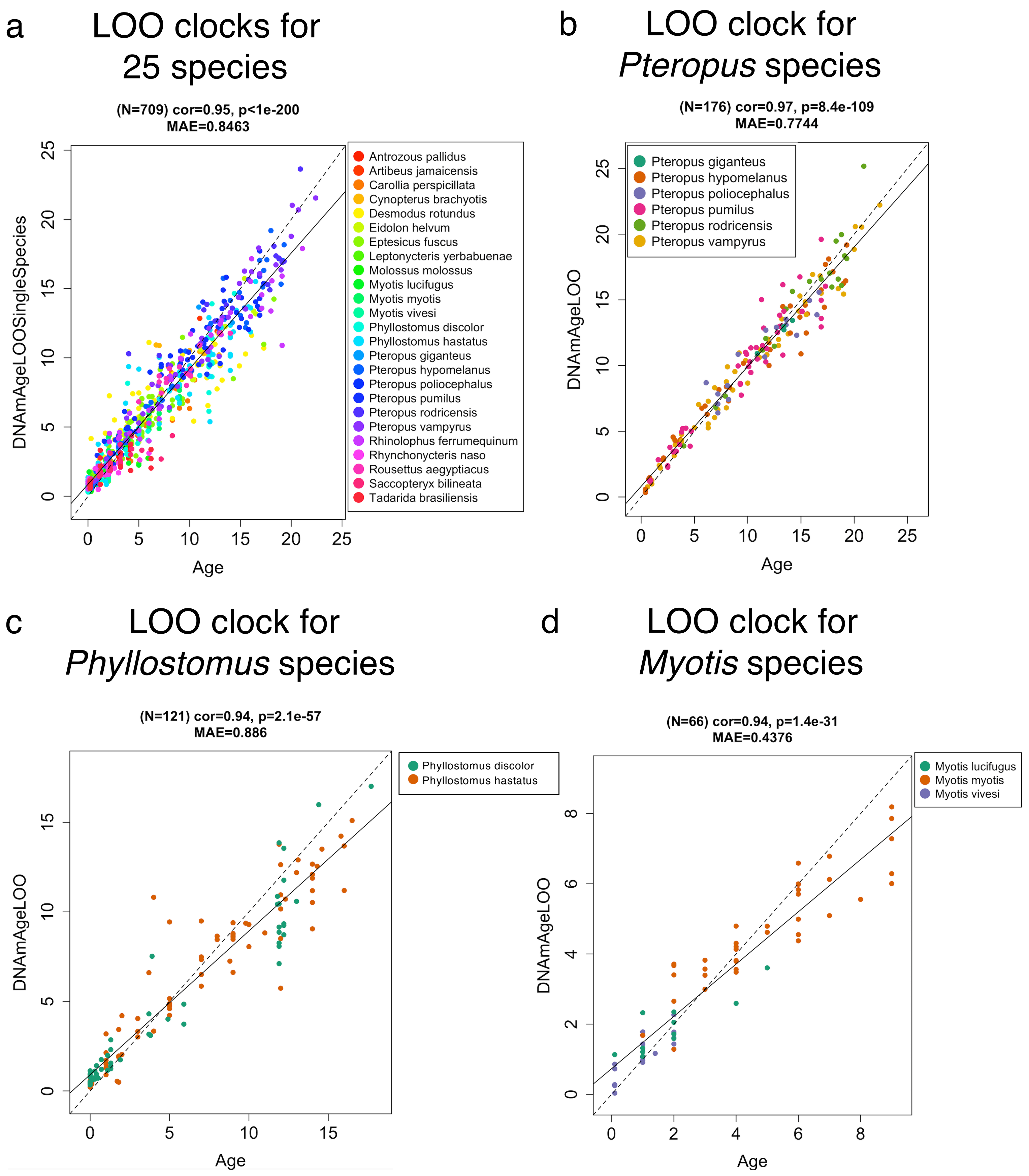
a
LOO clocks for 25 species
(N=709) cor=0.95, p<1e-200
MAE=0.8463
DNAmAgeLOOSingleSpecies
Age
Antrozous pallidus
Artibeus jamaicensis
Carollia perspicillata
Cynopterus brachyotis
Desmodus rotundus
Eidolon helvum
Eptesicus fuscus
Leptonycteris yerbabuenae
Molossus molossus
Myotis lucifugus
Myotis myotis
Myotis vivesi
Phyllostomus discolor
Phyllostomus hastatus
Pteropus giganteus
Pteropus hypomelanus
Pteropus poliocephalus
Pteropus pumilus
Pteropus rodricensis
Pteropus vampyrus
Rhinolophus ferrumequinum
Rhynchonycteris naso
Rousettus aegyptiacus
Saccopteryx bilineata
Tadarida brasiliensis
b
LOO clock for *Pteropus* species
(N=176) cor=0.97, p=8.4e-109
MAE=0.7744
DNAmAgeLOO
Age
Pteropus giganteus
Pteropus hypomelanus
Pteropus poliocephalus
Pteropus pumilus
Pteropus rodricensis
Pteropus vampyrus
c
LOO clock for *Phyllostomus* species
(N=121) cor=0.94, p=2.1e-57
MAE=0.886
DNAmAgeLOO
Age
Phyllostomus discolor
Phyllostomus hastatus
d
LOO clock for *Myotis* species
(N=66) cor=0.94, p=1.4e-31
MAE=0.4376
DNAmAgeLOO
Age
Myotis lucifugus
Myotis myotis
Myotis vivesi

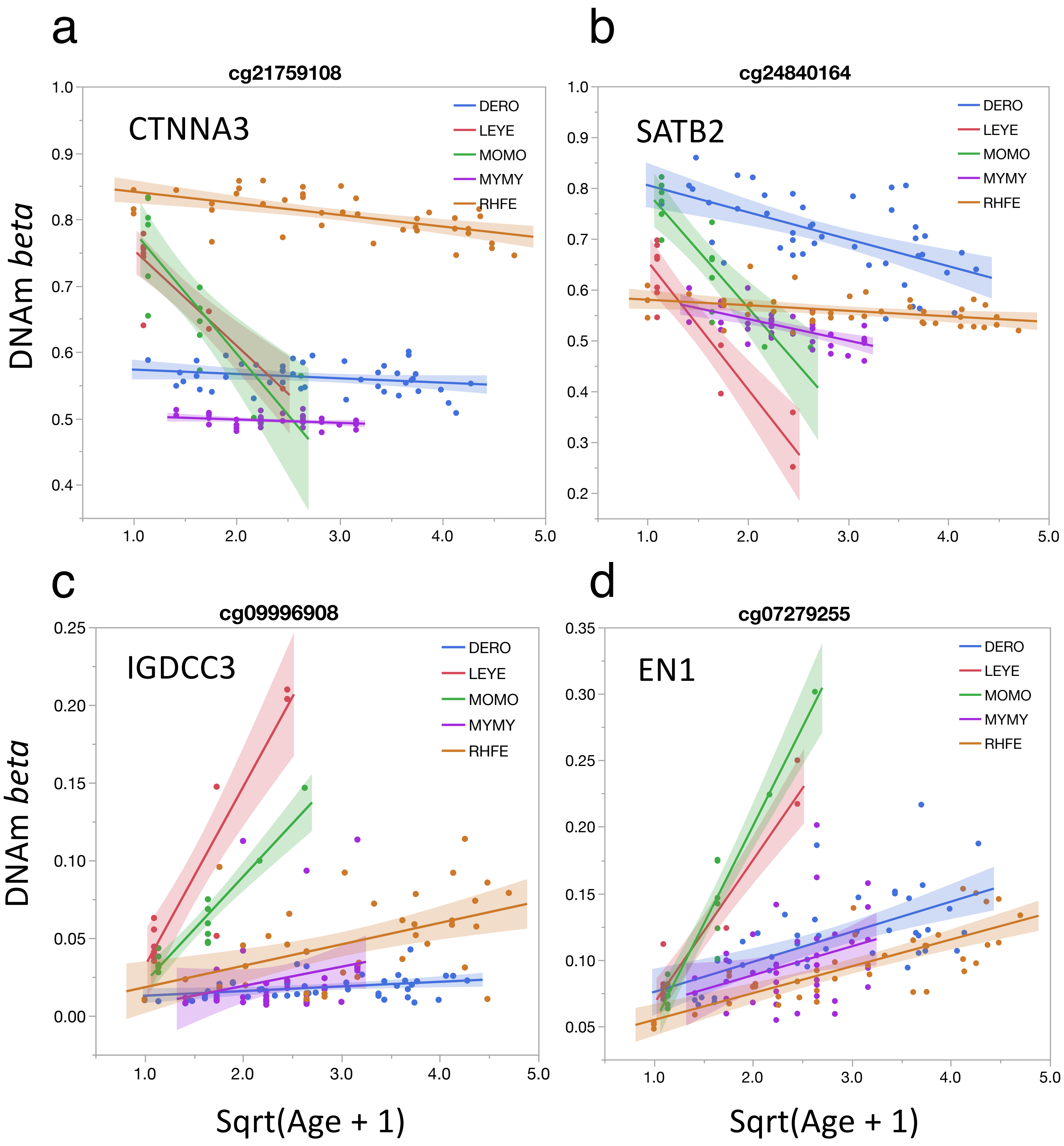
a
cg21759108
CTNNA3
b
cg24840164
SATB2
c
cg09996908
IGDCC3
d
cg07279255
EN1
DERO
LEYE
MOMO
MYMY
RHFE
DNAm beta
Sqrt(Age + 1)

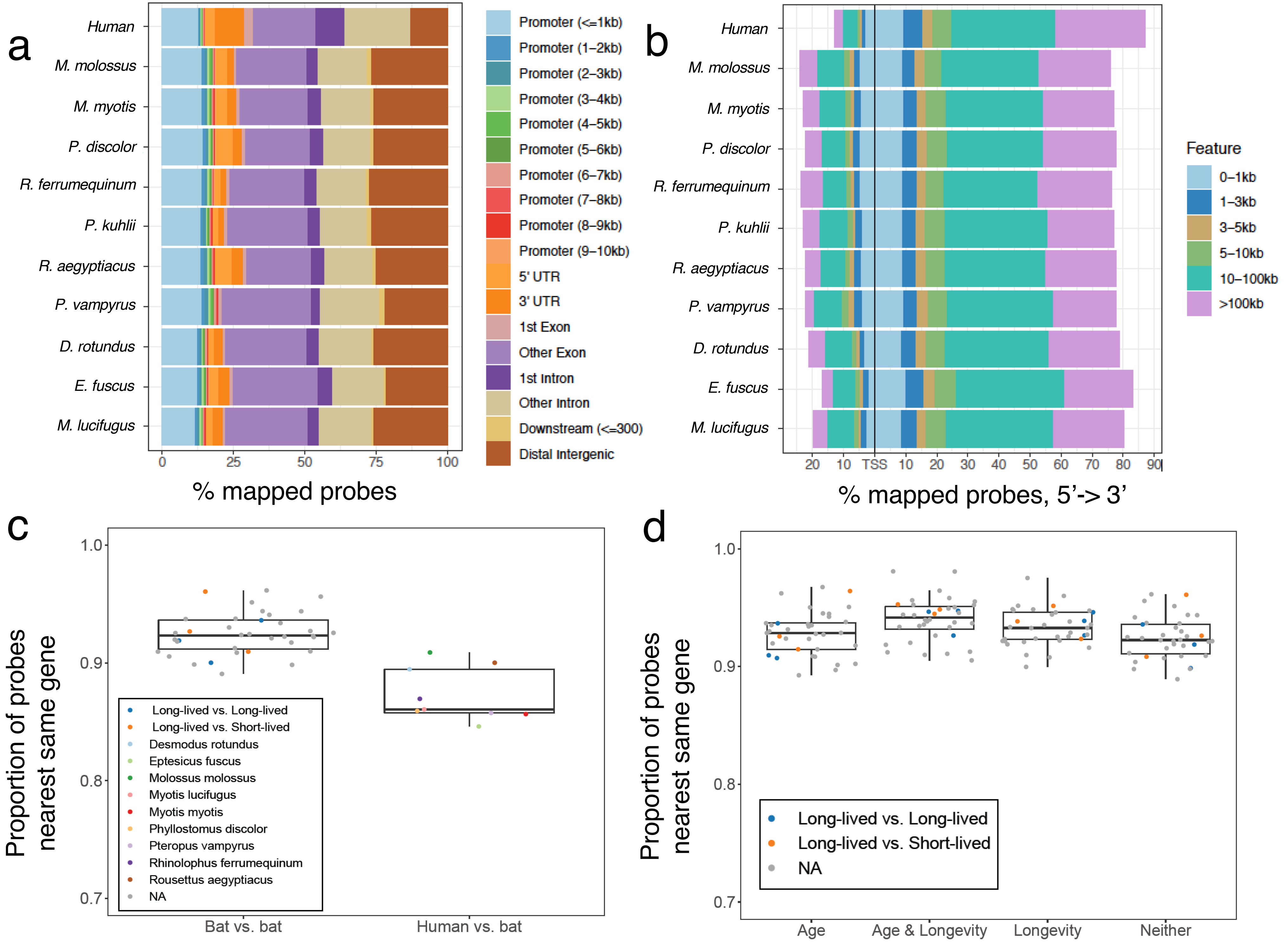

a
Human
M. molossus
M. myotis
P. discolor
R. ferrumequinum
P. kuhlii
R. aegyptiacus
P. vampyrus
D. rotundus
E. fuscus
M. lucifugus
0
25
50
75
100
% mapped probes
Promoter (<=1kb)
Promoter (1–2kb)
Promoter (2–3kb)
Promoter (3–4kb)
Promoter (4–5kb)
Promoter (5–6kb)
Promoter (6–7kb)
Promoter (7–8kb)
Promoter (8–9kb)
Promoter (9–10kb)
5' UTR
3' UTR
1st Exon
Other Exon
1st Intron
Other Intron
Downstream (<=300)
Distal Intergenic
b
20 10 TSS 10 20 30 40 50 60 70 80 90
% mapped probes, 5'-> 3'
Feature
0–1kb
1–3kb
3–5kb
5–10kb
10–100kb
>100kb
c
Proportion of probes nearest same gene
1.0
0.9
0.8
0.7
Bat vs. bat
Human vs. bat
Long-lived vs. Long-lived
Long-lived vs. Short-lived
Desmodus rotundus
Eptesicus fuscus
Molossus molossus
Myotis lucifugus
Myotis myotis
Phyllostomus discolor
Pteropus vampyrus
Rhinolophus ferrumequinum
Rousettus aegyptiacus
NA
d
Proportion of probes nearest same gene
1.0
0.9
0.8
0.7
Age
Age & Longevity
Longevity
Neither
Long-lived vs. Long-lived
Long-lived vs. Short-lived
NA

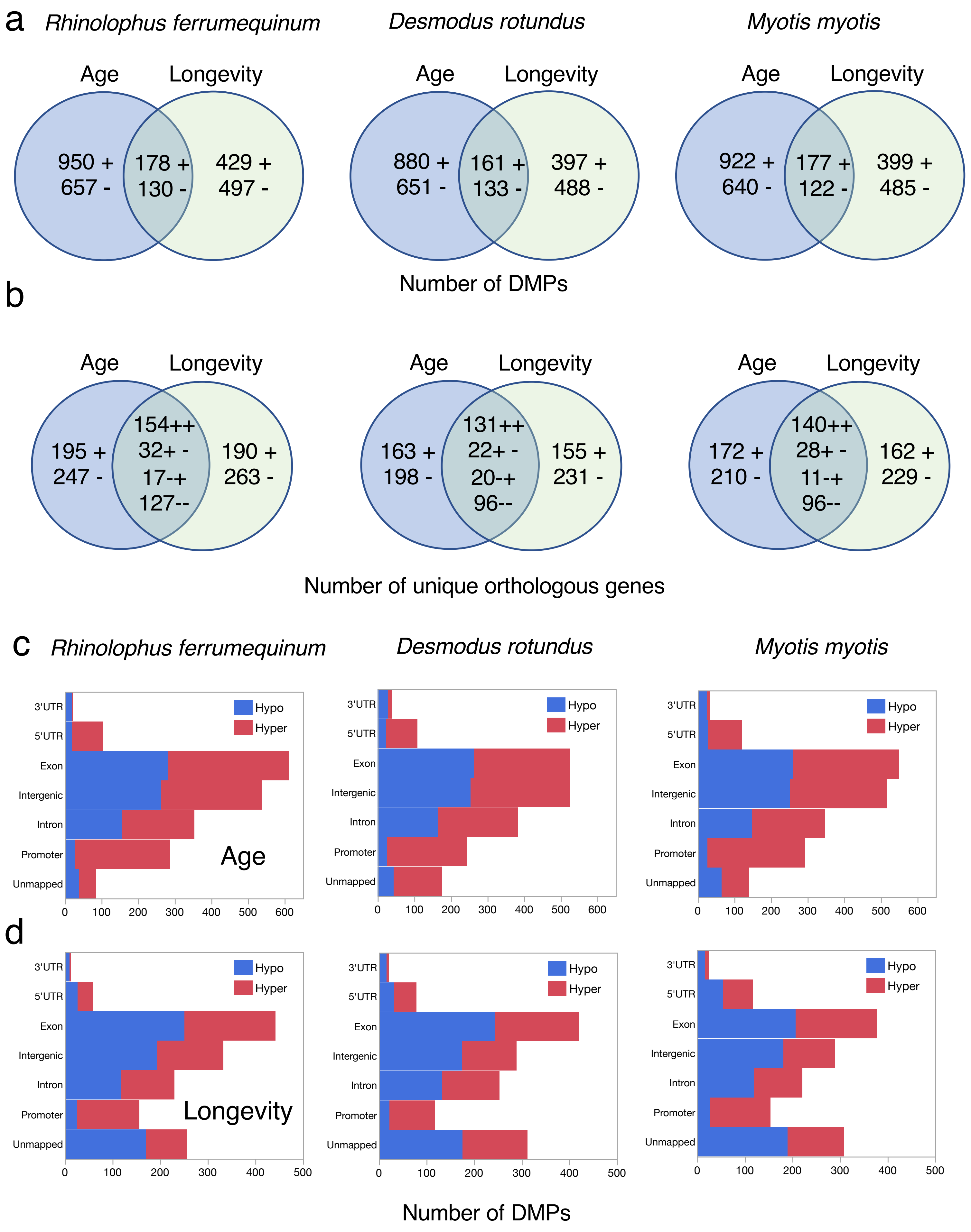
a
Rhinolophus ferrumequinum
Desmodus rotundus
Myotis myotis
Age
Longevity
950 +
657 -
178 +
130 -
429 +
497 -
880 +
651 -
161 +
133 -
397 +
488 -
922 +
640 -
177 +
122 -
399 +
485 -
Number of DMPs
b
195 +
247 -
154++
32+ -
17-+
127--
190 +
263 -
163 +
198 -
131++
22+ -
20-+
96--
155 +
231 -
172 +
210 -
140++
28+ -
11-+
96--
162 +
229 -
Number of unique orthologous genes
c
d
3'UTR
5'UTR
Exon
Intergenic
Intron
Promoter
Unmapped
Hypo
Hyper
Age
Longevity
Number of DMPs

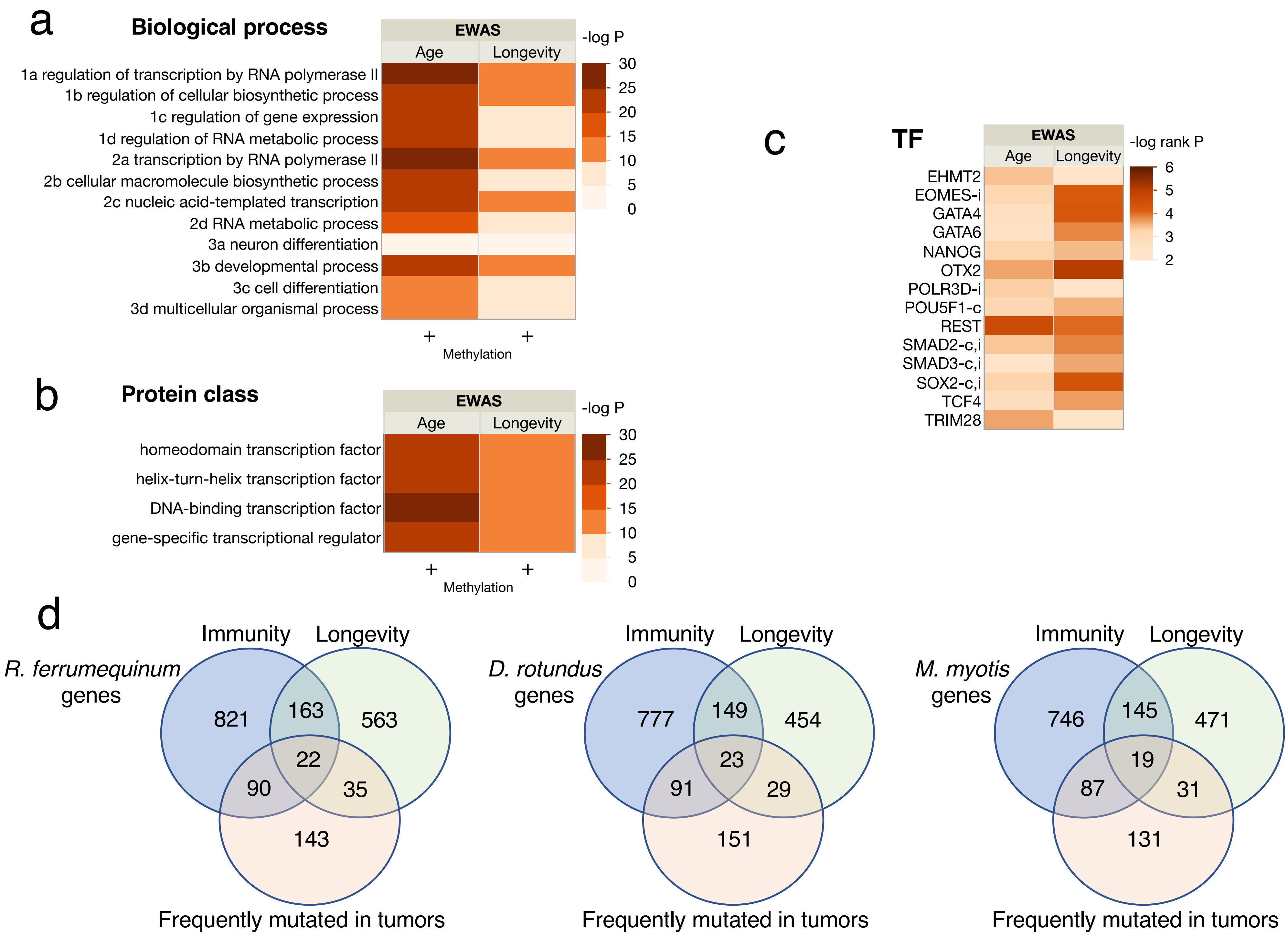

a
Biological process
EWAS
Age
Longevity
-log P
30
25
20
15
10
5
0
1a regulation of transcription by RNA polymerase II
1b regulation of cellular biosynthetic process
1c regulation of gene expression
1d regulation of RNA metabolic process
2a transcription by RNA polymerase II
2b cellular macromolecule biosynthetic process
2c nucleic acid-templated transcription
2d RNA metabolic process
3a neuron differentiation
3b developmental process
3c cell differentiation
3d multicellular organismal process
+
+
Methylation
b
Protein class
EWAS
Age
Longevity
-log P
30
25
20
15
10
5
0
homeodomain transcription factor
helix-turn-helix transcription factor
DNA-binding transcription factor
gene-specific transcriptional regulator
+
+
Methylation
c
TF
EWAS
Age
Longevity
-log rank P
6
5
4
3
2
EHMT2
EOMES-i
GATA4
GATA6
NANOG
OTX2
POLR3D-i
POU5F1-c
REST
SMAD2-c,i
SMAD3-c,i
SOX2-c,i
TCF4
TRIM28
d
Immunity
Longevity
R. ferrumequinum genes
821
163
563
22
90
35
143
Frequently mutated in tumors
Immunity
Longevity
D. rotundus genes
777
149
454
23
91
29
151
Frequently mutated in tumors
Immunity
Longevity
M. myotis genes
746
145
471
19
87
31
131
Frequently mutated in tumors
